# Rapid and accurate SNP genotyping of clonal bacterial pathogens with BioHansel

**DOI:** 10.1101/2020.01.10.902056

**Authors:** Geneviève Labbé, Peter Kruczkiewicz, Philip Mabon, James Robertson, Justin Schonfeld, Daniel Kein, Marisa A. Rankin, Matthew Gopez, Darian Hole, David Son, Natalie Knox, Chad R. Laing, Kyrylo Bessonov, Eduardo Taboada, Catherine Yoshida, Kim Ziebell, Anil Nichani, Roger P. Johnson, Gary Van Domselaar, John H.E. Nash

**Affiliations:** National Microbiology Laboratory, Public Health Agency of Canada, Canada; Department of Medical Microbiology & Infectious Diseases, University of Manitoba, Winnipeg, Manitoba, Canada

**Keywords:** *k*-mer, core SNP, canonical SNP, subtyping, fast WGS data analysis, bioinformatics

## Abstract

BioHansel performs high-resolution genotyping of bacterial isolates by identifying phylogenetically informative single nucleotide polymorphisms (SNPs), also known as canonical SNPs, in whole genome sequencing (WGS) data. The application uses a fast *k*-mer matching algorithm to map pathogen WGS data to canonical SNPs contained in hierarchically structured schemas and assigns genotypes based on the detected SNP profile. Using modest computing resources, BioHansel efficiently types isolates from raw sequence reads or assembled contigs in a matter of seconds, making it attractive for use by public health, food safety, environmental, and agricultural authorities that wish to apply WGS methodologies for their surveillance, diagnostics, and research programs. BioHansel currently provides canonical SNP genotyping schemas for four prevalent *Salmonella* serovars—Typhi, Typhimurium, Enteritidis and Heidelberg—as well as a schema for *Mycobacterium tuberculosis*. Users can also supply their own schemas for genotyping other organisms. BioHansel’s quality assurance system assesses the validity of the genotyping results and can identify low quality data, contaminated datasets, and misidentified organisms. BioHansel is targeted to support surveillance, source attribution, risk assessment, diagnostics, and rapid screening for public health purposes, such as product recalls. BioHansel is an open source application with packages available for PyPI, Conda, and the Galaxy workflow manager. In summary, BioHansel performs efficient, rapid, accurate, and high-resolution classification of bacterial genomes from sequence reads or assembled contigs on standard computing hardware. BioHansel is suitable for use as a general research tool as well as in fully operationalized WGS workflows at the front lines of infectious disease surveillance, diagnostics, and outbreak investigation and response.

**Impact statement:** Public health, food safety, environmental, and agricultural authorities are currently engaged in a global effort to incorporate whole genome sequencing technologies into their infectious disease research, surveillance, and outbreak investigation programs. Its widespread adoption, however, has been impeded by two major obstacles: the need for high performance computing to generate results and the expert knowledge required to interpret and communicate those results. BioHansel addresses these limitations by rapidly genotyping pathogens from whole genome sequence data in an accurate, simple, familiar, and easily sharable manner using standard computing resources. BioHansel provides a compact and readily interpretable genotype based on canonical SNP genotyping schemas. BioHansel’s genotyping nomenclature encodes the pathogen’s position in its population structure, which simplifies and facilitates its comparison with actively circulating strains and historical strains. The genotyping information provided by BioHansel can identify points of intervention to prevent the spread of pathogenic bacteria, screen for the presence of priority pathogens, and perform source attribution and risk assessment. Thus, BioHansel serves as a readily accessible and powerful WGS method, implementable on a laptop, for genotyping pathogens to detect, monitor, and control the emergence and spread of infectious disease through surveillance, screening, diagnostics, and outbreak investigation and response activities.

**Data summary:** BioHansel is a Python 3 application available as PyPI, Conda Galaxy Tool Shed packages. It is an open source application distributed under the Apache License, Version 2.0. Source code is available at https://github.com/phac-nml/biohansel. The BioHansel user guide is available at https://bio-hansel.readthedocs.io/en/readthedocs/. Supplementary Materials are available at https://github.com/phac-nml/biohansel-manuscript-supplementary-data.

The authors confirm all supporting data, code and protocols have been provided within the article or through supplementary data files.

## Introduction

Public health, animal health, food safety, and environmental authorities around the world are actively working to operationalize whole genome sequencing (WGS) technologies for their infectious disease diagnostics, surveillance, and outbreak detection and response programs. Data analysis is the biggest challenge facing the adoption of WGS for these applications (Deurenberg et al., 2017; Nadon et al., 2017). The genomic data analysis pipelines that serve these mission-critical programs must implement robust, reproducible, and computationally tractable analytical methods that generate accurate and informative results (PHG Foundation 2015). Data sharing is another challenge, especially the sharing of pathogen WGS reads, due to outmoded data governance policies, insufficient network bandwidth, and scant infrastructure to facilitate its distribution among stakeholders in different institutions or in different health jurisdictions. Subtyping data derived from pathogen WGS data currently serves as a better alternative to WGS for communicating infectious disease information, since the sharing of subtyping data is already a well-established and long-standing practice among health authorities and is compatible with the existing policies that govern the sharing of such data (European Centre for Disease Prevention and Control, 2015). Subtype data is also much more compact than WGS data and requires negligible network bandwidth to transmit).

The ability of WGS to resolve bacterial relationships is unmatched by any other molecular technology, and its adoption by public health and food safety authorities has revolutionised our ability to track and respond to disease outbreaks (Nadon *et al.*, 2017; Tang *et al.*, 2017; Besser *et al.*, 2018; Zakham *et al.*, 2019). WGS is especially useful for highly clonal organisms, since there are typically only a small number of genetic changes that distinguish a clonal disease outbreak from unrelated contemporaneously circulating strains, and the genetic changes that provide this discrimination are important to contextualize, monitor, and respond to such outbreaks. Clonally acquired genetic changes manifest predominantly as point mutations, such as single nucleotide polymorphisms (SNPs) and small insertions or deletions (indels). Additional non-clonal sources of genetic diversity include recombination and the gain or loss of mobile elements such as transposons, bacteriophage, and plasmids. Several approaches exist that exploit this genomic diversity to estimate the genetic and epidemiological relatedness of infectious diseases, the most popular being the gene-by-gene methods and the SNP-based methods (Schürch *et al.*, 2018). The choice of typing approach depends on its intended application (for a review see Nadon *et al.*, 2017), although in general, gene-by-gene approaches are popular for routine surveillance activities, whereas SNP-based approaches are used when extra discriminatory power is required, or for pathogens that are not amenable to gene-by-gene methods. Comparisons of gene-by-gene and SNP-based approaches show that these methods both produce concordant and complementary results (Katz *et al.*, 2017; Whaley *et al.*, 2018; Jagadeesan *et al.*, 2019). Regardless of the method used, the analysis of WGS data can be highly complex and can require high performance computing and specialized genomics expertise to carry out and to interpret the results (Carriço *et al.*, 2018).

Gene-by-gene methods estimate genetic relatedness by indexing the allelic diversity of a designated set of genes. Each allelic variant is assigned a unique identifier and the genetic distance between two isolates is computed as the number of allelic differences that exist between them (Maiden *et al.*, 2013). The core genome multi-locus sequence typing (cgMLST) approach considers only the genes that are common to all (or nearly all) genomes for a given pathogen, whereas the whole-genome multi-locus sequence typing (wgMLST) approach considers both core and accessory genes. Gene-by-gene approaches generate allelic profiles that can be assigned to a categorical type; in this way, a large amount of genomic data is collapsed into a simple and highly compact form which facilitates pathogen genomic data sharing. They do not, however, consider the individual nucleotide variants that distinguish the alleles, nor do they consider polymorphisms occurring in intergenic regions (Jagadeesan *et al.*, 2019). While this approach helps to ameliorate the confounding effects of recombination on genotype assignment, it renders the gene-by-gene approaches somewhat less discriminatory than their SNP-based counterparts. Gene-by-gene methods can also be sensitive to contamination and mixed colonies, although this problem extends to all genotyping approaches (Page *et al.*, 2017). Several pipelines are available for gene-by-gene analysis of WGS data; popular examples include Ridom SeqSphere+ (Jünemann *et al.*, 2013), BIGSdb (Jolley and Maiden, 2010; Jolley *et al.*, 2018), BioNumerics (Applied Maths), EnteroBase (Alikhan *et al.*, 2018), and SRST2 (Inouye *et al.*, 2014).

SNP-based phylogenomic approaches are a popular alternative to gene-by-gene approaches. They typically rely on the selection of a high-quality reference genome against which all other isolates are compared, followed by the identification of the nucleotide variants that differentiate the reference and the query genomes. These approaches generate highly accurate phylogenies with the power—in theory—to distinguish isolates differing by a single base pair; however, a number of technical barriers exist that impede their routine application for infectious disease surveillance, outbreak detection, and response. One major barrier concerns the nature of the output—a phylogenetic tree—which provides context on the evolutionary relationship of a collection of isolates, but must be recomputed and reinterpreted as new isolate genomes become available. Traditional SNP methods do not assign genotype codes to the isolates (SnapperDB (Dallman *et al.*, 2018) is a notable exception); this frustrates surveillance and outbreak investigations since the lack of genotype offers no straightforward means for comparing actively circulating strains with historical strains or strains circulating in other health jurisdictions. Another quite prohibitive barrier is the high computing cost required to carry out the analysis, often requiring high performance computing to generate results on a timescale useful for outbreak investigations. Despite these barriers, SNP-based phylogenomic analysis pipelines are used extensively for outbreak investigation and response. Notable examples of SNP-based phylogenomic pipelines include the CFSAN SNP Pipeline (Davis *et al.*, 2015) used in the international GenomeTrakr network for detection of foodborne pathogens, the Lyve-SET hqSNP pipeline (Katz *et al.*, 2017) used for the analysis of enteric organisms at the U.S. Centers for Disease Control, the SNVPhyl pipeline (Petkau *et al.*, 2017) used for bacterial phylogenomics by the Canadian Public Health Laboratory Network, and the SnapperDB pipeline (Dallman *et al.*, 2018) used for national surveillance of enteric pathogens at Public Health England. Each of these pipelines has its own set of complex dependencies and assumptions that can influence the results (Petkau *et al.*, 2017; Katz *et al.*, 2017; Lynch *et al.*, 2016). In a recent review of 41 different SNP detection pipelines, the authors demonstrated that the choice of reference genome for SNP calling had a substantial effect on the variants identified, which can influence the phylogeny and thus its scientific and epidemiological interpretation (Bush *et al.*, 2019). SNP detection is sensitive also to recombination, sequence coverage, and the presence of contamination (Petkau *et al.*, 2017), although these sensitivities hold true for gene-by-gene methods as well.

One method for circumventing the difficulties inherent in traditional SNP-based workflows is to pre-identify the SNPs contained in the target organism’s core genome that capture the clonal evolutionary history of that organism. These phylogenetically informative SNPs, termed *canonical SNPs*, can be organized into hierarchically structured genotyping schemas. Like their gene-by-gene-based counterparts, canonical SNP-based approaches can generate compact, informative genotypes and have proven useful for source tracking, surveillance, trend analysis, and epidemiological investigations (Coll *et al.*, 2014; Wong *et al.*, 2016). Hierarchical SNP-based genotyping schemas have been designed for several organisms, including *M. tuberculosis* (Coll *et al.*, 2014), *Salmonella* Typhi (Wong *et al.*, 2016), *S.* Heidelberg (Labbe *et al.*, 2019) and *Bordetella pertussis* (van Gent *et al.*, 2011). Although promising, with the exception of *S.* Typhi and *M. tuberculosis*, the utility of these genotyping schemas—and hence the canonical SNP approach in general—has been limited by the lack of companion software to extract the informative SNPs and assign a genotype (Coll *et al.*, 2015; Khol *et al.*, 2018; Wong *et al.*, 2016).

Regardless of the chosen approach, an important and frequently encountered concern when using WGS for microbial genomic applications is the potential to encounter contamination, which can confound subtype assignment, and can be troublesome to detect. Mixed infections, which present similarly to contamination, can be similarly troublesome to detect (Khol *et al.*, 2018; Anyansi *et al.*, 2019). The most common sources of contamination include improperly isolated genomic material, environmental contamination, the libraries used to prepare the genomic material for sequencing, and DNA barcode “cross-talk” generated during the sequencing step (Wright and Vetsigian, 2016; Pettengill *et al.*, 2016; Goig *et al.*, 2019). To account for this complexity, contamination detection pipelines are commonly applied to newly generated WGS data prior to analysis. These pipelines normally assign reads to taxa using fast phylotyping tools such as Kraken (Wood and Salzberg, 2014), Centrifuge (Kim *et al.*, 2016), or Kaiju (Menzel *et al.*, 2016). However, these tools cannot reliably assign taxa beyond the species level and thus are unsuitable for identifying genomic contamination from the same species (Ye *et al.*, 2019; Sczyrba *et al.*, 2017; McIntyre *et al.*, 2017). Few options exist for identifying intra-species contamination in highly clonal microbial genomic sequence data (Anyansi *et al.*, 2019). ConFindr identifies intra-species contamination by searching for the presence of multiple variant bases in a set of conserved single-copy ribosomal protein-encoding genes (Low *et al.*, 2019). However, this approach is unable to detect contamination from highly clonal lineages that may have little to no diversity at these loci. QuantTB (Anyansi *et al.*, 2019) identifies mixed infections by comparing the SNP profiles of *M. tuberculosis* samples to those present in a curated reference database of genome sequences (Anyansi *et al.*, 2019). The VCFMIX program (Wyllie *et al.*, 2018) similarly maps reads to a reference genome followed by statistical analysis of detected SNPs for known genotypes. A simple modification of the QuantTB and VCFMIX approaches to identify contamination in clonal populations of infectious disease is to compare against lineage-specific canonical SNPs targeted by *k*-mers, eliminating the need for whole genome mapping.

As health authorities increasingly adopt WGS for microbial surveillance, diagnostics, and outbreak response, tools that provide rapid, accurate results with minimal computing requirements and generate results in a useful, familiar, and easily sharable form are in high demand. We developed BioHansel to meet this demand. BioHansel uses a fast *k*-mer mapping approach to identify canonical SNPs in pathogen WGS data and uses these SNPs to assign high-resolution genotypes that encode the isolate’s position in a population structure. Here we describe BioHansel’s design and demonstrate its ability to rapidly and accurately genotype isolates for *S*. Heidelberg, *S*. Enteritidis, *S*. Typhimurium, *S*. Typhi, and *M. tuberculosis* using modest computing resources. We then demonstrate BioHansel’s intrinsic ability to assess input genomic data for the presence of intra-species contamination. We also demonstrate BioHansel’s memory efficiency, compute speed, and scalability by performing a benchmarking study against Snippy, a popular traditional microbial SNP variant detection pipeline (Seemann *et al.*, 2015).

## Methods

### Canonical SNP genotyping schemas

Most SNP-based genotyping schemas use canonical SNPs to assign genotypes. We designed our BioHansel schemas by conducting a deep manual investigation of the population structure for each pathogen, culminating in the identification of canonical SNPs that hierarchically partition that pathogen’s population structure. Each schema consists of a set of canonical SNPs, their genomic context, and a hierarchically encoded nomenclature that records the SNP’s position in the hierarchy. Schemas are constructed from these SNPs by specifying, for each SNP, an “inclusion” SNP variant and an optional “exclusion” SNP variant that defines how the SNP partitions the population structure into subpopulations, as detailed below.

We provide schemas for *S*. Enteritidis (SE), *S*. Typhimurium (ST), *S*. Heidelberg (SH), *M*. *tuberculosis* (MTB), and *S*. Typhi (Typhi). Users can also supply BioHansel with their own custom schemas, which are encoded in a simple FASTA format. The SE, ST, and SH schemas were developed at the Public Health Agency of Canada’s National Microbiology Laboratory; the MTB and Typhi schemas were adapted from previously published genotyping schemas developed by Coll *et al.* (2014) and Wong *et al.* (2016), respectively. The SE and ST schemas were developed and tested using > 20,000 publicly available WGS datasets drawn from all available SE and ST WGS datasets available in the NCBI SRA database at the time of schema construction and thus represent international sources. The SH schema was developed from > 2,000 isolates drawn from mostly Canadian and American sources. Details of the schema construction process along with all five schemas, the reference genome sequences used to construct the schemas, the phylogenies constructed from the canonical SNPs, and the genotypes along with their associated SNP profile variants are provided in the Supplementary Materials (https://github.com/phac-nml/biohansel-manuscript-supplementary-data).

#### BioHansel schema design

We designed BioHansel to work with hierarchical genotyping schemas that follow the lineage genotyping framework developed for *M. tuberculosis* (Coll *et al.*, 2014). This approach provides a flexible, adaptable, and extensible framework for representing clonal population structures. We provide a brief synopsis of the BioHansel schema creation procedure here; a more detailed procedure is available in the Supplementary Materials.

The population structure for a given organism is hierarchically clustered into lineages, clades, subclades, and so on. Genotypes are assigned to every node in the population structure using a dotted-decimal notation (*X*_1_, *X*_1_. *X*_2_, *X*_1_. *X*_2_. *X*_3_, *etc*.) that encodes the node’s position in the hierarchy. For quality control purposes, one or more unique inclusion SNP *k*-mers must support each labelled genotype in the lineage. Each inclusion *k*-mer consists of the SNP variant that distinguishes a clade from the rest of the population along with equal-length flanking sequences that uniquely specify that SNP in the target population. (We found that a *k*-mer length of 33 nucleotides demonstrated good sensitivity and specificity for our schemas, but BioHansel can use *k*-mers of any size, including variable size *k*-mers.) Only dichotomous, core genome SNPs (*i.e.*, only a single variant form found throughout the whole population) were considered as candidate canonical SNPs. This excluded SNPs present only in a subpopulation (*i.e.*, those contained in indel regions) from our genotyping schemas. BioHansel can, however, accept schemas with SNPs in indel regions, and it also accepts *k*-mers containing degenerate bases.

BioHansel requires only that an inclusion *k*-mer be specified for each SNP defining a subpopulation. If the optional exclusion *k*-mer is additionally specified, then it will be used for quality assessment and quality control (see the BioHansel Read-the-Docs for more detail). The addition of exclusion *k*-mers ensures that all the expected SNP targets are present in the dataset. A large number of missing targets can indicate poor quality sequence data, inadequate genome coverage, or recombination. The presence of both inclusion and exclusion *k*-mers for a given SNP position indicates possible cross-contamination or mixed culture. BioHansel issues warnings when it encounters these conditions.

### BioHansel

BioHansel accepts pathogen genome sequence reads or assembled contigs along with a corresponding canonical SNP genotyping schema. BioHansel forms quality assessment and (QA) and quality control (QC) on the supplied sequence data, searches the data for matching *k*-mers contained in the schema, and assigns a genotype based on the detected SNP profile. We summarize this workflow in Figure 1.

**Figure 1.**
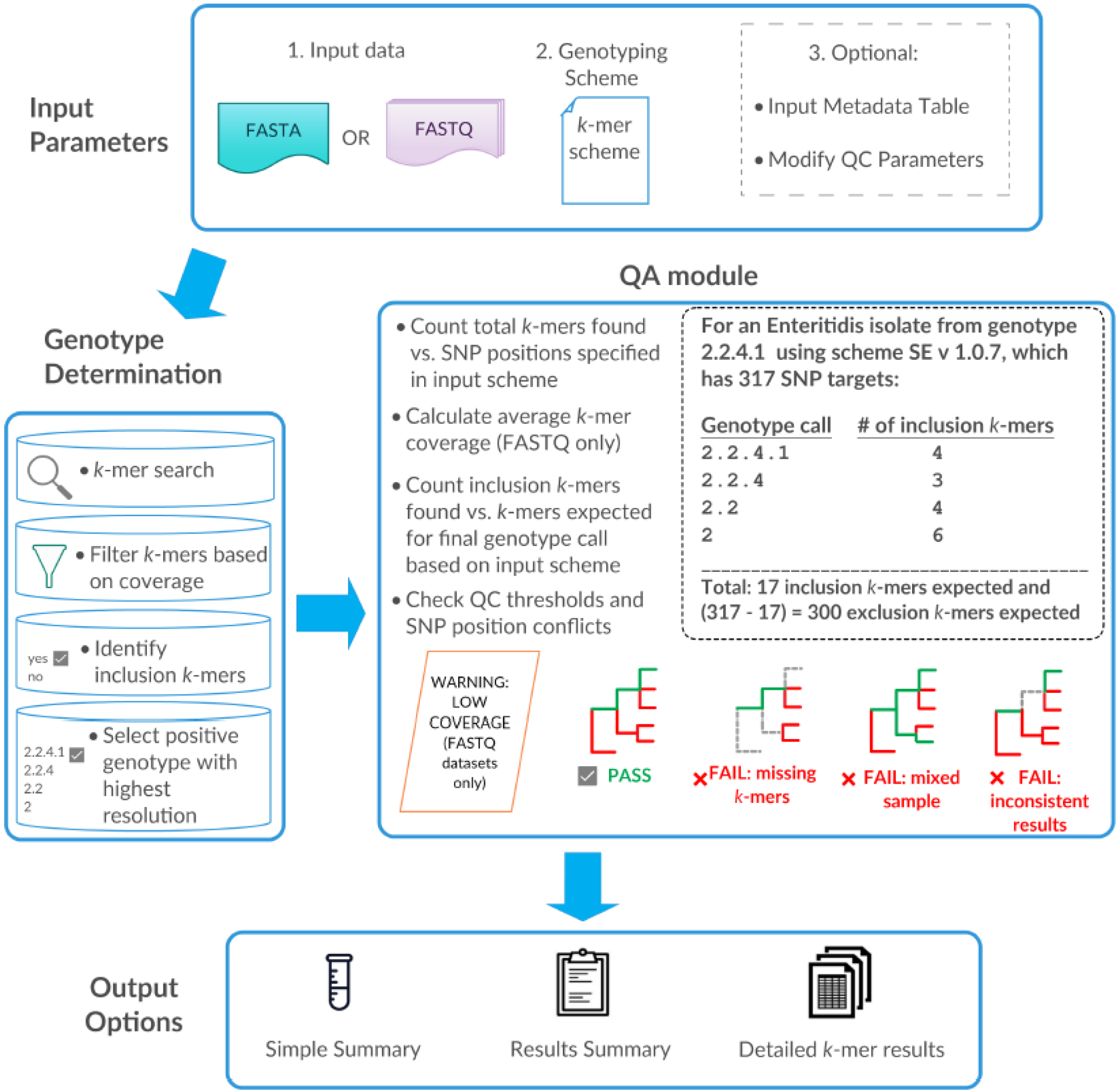
BioHansel workflow. BioHansel accepts raw reads or genome assemblies as input, along with a corresponding genotyping schema. The tool uses *k*-mers in the genotyping schema to find canonical SNPs in the pathogen sequence data and assign the highest-resolution genotype specified by the matching SNP profile. BioHansel performs quality assessment by computing the detected SNP coverage (for read data), checking the consistency of the detected SNPs with the population structure defined in the SNP schema, and checking for possible contamination. BioHansel then creates one or more output result files as specified by the user; examples of each output file type are provided in Supplementary Table S2.

#### SNP detection algorithm

BioHansel searches for *k*-mers within the input sequence data using the Aho–Corasick algorithm (Aho and Corasick, 1975) implemented in the *pyahocorasick* Python library (Muła, 2019). The Aho–Corasick algorithm rapidly and efficiently searches for the supplied set of reads or assembled contigs for *k*-mers defined in the corresponding schema. BioHansel expands the *k*-mers containing degenerate bases to include all possible variants and searches the input data for these variants.

#### Quality Assessment and Quality Control

BioHansel’s QA/QC module provides insight into the validity of obtained genotyping results by reporting the consistency of the detected SNPs with the population structure defined in the associated schema (Figure 1). Several features are assessed, including the total number of *k*-mer targets identified, the presence of a single SNP variant for a given genotype, the phylogenetic consistency of detected SNP profile, and the depth of coverage for detected SNPs (when read data is provided as input; see below). When both inclusion and exclusion *k*-mers are provided, BioHansel tallies the number of *k*-mer targets found in the genomic data to assess the quality of the results: isolates that are missing a large number of target SNPs indicate poor quality sequence data, inadequate read sequence coverage, or possible recombination. If BioHansel is provided with read data, it will calculate the coverage depth of each detected SNP, report the average coverage across all detected SNPs, and issue a warning if it falls below the minimum coverage threshold. BioHansel determines intermediate genotypes or ambiguous genotyping results by examining the phylogenetic consistency of the detected SNP profile. Phylogenetic consistency is determined by selecting the genotype(s) with the highest resolution within the hierarchy and then counting the number of detected SNPs present in the lower hierarchical levels that support the final genotype.

BioHansel includes checks for confidence and ambiguity. The tool assigns as *ambiguous* genotyping results that are missing ≥ 3 target SNPs for a given genotype. When both the inclusion and exclusion *k*-mers are provided in the schema, BioHansel reports a *fail* status if it cannot identify the exclusion *k*-mer targets contained in the subpopulation defined by the highest-resolution genotype identified in the supplied sequence data. If all of the inclusion *k*-mer SNP targets from a single level of the hierarchy are missing, the isolate is assigned a status of *inconclusive*. BioHansel also detects heterozygosity at a variant site if it identifies both inclusion and exclusion *k*-mer targets in the WGS data; this can be indicative of an intra-strain mixture or contamination.

#### Metadata Table

BioHansel accepts an optional metadata table that can enhance its genotyping report with additional contextual information. We include metadata tables for the Typhi and MTB schemas that were adapted for BioHansel from previously published work (Wong *et al.*, 2016; Coll *et al.*, 2014) (see Supplementary Table S3). These metadata tables supplement BioHansel’s genotype assignment with the corresponding assignment from the original publications. The metadata fields are arbitrary, and users can modify their schemas with as many metadata fields as they like; for example, known source associations, associated outbreak events, geographical locations, collection date, disease severity, *etc*.

### Analysis of retrospective outbreaks with BioHansel

We validated BioHansel using genome sequence reads from three retrospective Canadian outbreaks of *S*. Heidelberg (Bekal *et al.*, 2016) previously analysed using the SNVPhyl phylogenomics pipeline (Petkau *et al.*, 2017). We used SNVPhyl with the *S.* Heidelberg strain SL476 reference genome to identify high quality SNPs from 59 *S*. Heidelberg isolates. The list of accessions for these isolates along with their BioHansel genotype codes are provided in Supplementary Table S4. Phylogenetic trees were built with Mega7 (Kumar *et al.*, 2016) and were visualized with iTOL (Letunic and Bork, 2019). We also analysed the same isolates by wgMLST using BioNumerics v. 7.6.3 (Applied Maths, Sint-Martens-Latem, Belgium) using the *Salmonella* wgMLST scheme with 4,396 alleles. Phylogenies were generated from the detected allelic profiles using unweighted pair group method with arithmetic mean (UPGMA).

### BioHansel SNP detection validation

BioHansel’s canonical SNP detection algorithm was validated against Snippy (Seemann, 2015), a popular traditional SNP-calling workflow for microbial genomes. BioHansel’s genotype assignments for the ST genotyping schema were compared against the results from Wong *et al.* (Wong *et al.*, 2016). We validated the concordance of BioHansel genotyping results with Snippy using a combination of real and synthetic data sets. Detailed parameter settings for all programs used for validation are provided in Supplementary Worksheet 1.

#### BioHansel and Snippy SNP-detection concordance for *S.* Typhi

We validated BioHansel’s genotype assignments for *S*. Typhi with randomly selected genome assemblies drawn from the Enterobase public repository of > 7,200 *S*. Typhi genome sequences (Alikhan *et al.*, 2018). The selected assemblies were serotyped using SISTR, a Salmonella *in silico* serotyping tool (Yoshida *et al.*, 2016), to confirm their *S*. Typhi serotype. This initial dataset was filtered to include only sequences with ≥ 40X coverage and a genome assembly size between 4.4 Mb and 5.5 Mb. To maximize the diversity of isolates, only a single representative isolate sequence was chosen for each sequence type assigned by Enterobase. After filtering and dereplication, 1,000 isolates were selected and their reads were downloaded for validation (Supplementary Table S5).

BioHansel v. 2.2.0 and Snippy v. 4.3 were run on 1,000 sets of *S*. Typhi sequence reads, and their SNP calling concordance was assessed (Table 1). BioHansel does not call SNPs in the same sense that a traditional SNP calling workflow does; instead, it reports on whether or not an exact match to the *k*-mer targeting that SNP was found in the sequence read data. In order to compare Snippy and BioHansel, we considered only the 68 canonical SNPs contained in the *S.* Typhi schema. For both BioHansel and Snippy, the number of assembled reads or *k*-mers supporting the presence of each SNP was tabulated, and the SNP variant supported by the majority of the reads was identified. Each position was assigned one of three states for each tool: *unambiguous* (a single SNP variant detected), *mixed* (both SNP variants detected) or *null* (missing/absent). Python scripts and a Nextflow workflow were written to extract the SNP variants identified by BioHansel and Snippy (Supplementary Materials). The effect of SNP-coverage cut offs on SNP detection ability were examined using 3X, 6X, and 8X coverage cut offs to simulate the level of experimental error found at the range of total coverages typically targeted for these pathogens when sequenced on the Illumina MiSeq platform (i.e., > 40X). The mean mapping coverage for each read set was was calculated from the Snippy BAM files with "samtools depth” (Li *et al.*, 2009).

**Table 1:** Genotyping results from BioHansel and Snippy using 68 SNP targets with 1,000 *S*. Typhi genomes at three coverage thresholds (*n* = 68,000 comparisons at each genome coverage level).

| Minimum Coverage Cut Off | Calls Agree | Discordant Majority Calls | Missed Calls, Both | Mixed Calls, Both | Missed Calls, Snippy | Mixed Calls, Snippy | Missed Calls, BioHansel | Mixed Calls, BioHansel |
| --- | --- | --- | --- | --- | --- | --- | --- | --- |
| 3X | 67,912 | 0 | 22 | 43 | 22 | 92 | 32 | 72 |
| 6X | 67,975 | 0 | 23 | 5 | 23 | 19 | 34 | 5 |
| 8X | 67,985 | 0 | 24 | 0 | 24 | 4 | 35 | 0 |

#### Validation of BioHansel genotype assignment against published genotypes

The genotyping results produced by BioHansel v. 2.2.0 were compared with the genotyping results produced by Genotyphi (Holt, 2019; Wong *et al.*, 2016, Britto *et al.*, 2018, Rahman *et al.*, 2019) on the same set of 1,910 *S*. Typhi isolates used in the original publication (Wong *et al.*, 2016). BioHansel’s Typhi schema v. 1.2.0 was adapted from the SNPs identified by Wong *et al.* to include the 16 bases flanking each SNP in the reference genome CT18, for a total *k*-mer length of 33 bases (Wong *et al.*, 2016). Both an inclusion and an exclusion *k*-mer was specified for each SNP location based on assignments by Wong *et al.* The nomenclature was adjusted to conform to the strict hierarchical format expected by BioHansel.

### Contamination Detection

#### BioHansel and Snippy SNP-calling concordance on artificial contamination datasets

In order to compare the mixed base-calling results between BioHansel and Snippy, we chose to maximize the number of positions that would yield an ambiguous base call by selecting two isolate sequences from the two most divergent genotypes in each of our five genotyping schemas (accession numbers are provided in Supplementary Table S6). Each sequence was assembled using Unicycler v. 0.47 (Wick *et al.*, 2017) using the default settings. We used ART Illumina v. 3.11.14 (Huang *et al.*, 2012) to generate from each of these sequences a synthetic paired-end Illumina read dataset with 500X coverage. We used the *sample* command from Seqtk v. 1.3 (Li, 2019) with a fixed seed of 100 to randomly subsample the reads from these datasets at various proportions and then merged them to generate our final contamination datasets with simulated contamination levels and coverages of 0% (0X), 1.67% (2X), 5% (6X), 10% (12X), 13.33% (16X), 16.67% (20X), 33.33% (40X), and 50% (60X), with a total coverage of 120X for each mixed pair (see list of commands in Supplementary Worksheet 2).

BioHansel v. 2.2.0 and Snippy v. 4.3 were run with their default settings on the artificial contamination datasets, and their SNP calling concordance was assessed for each SNP target in each of BioHansel’s five genotyping schemas. The contaminated datasets were analysed by Snippy with the reference sequences used to generate the genotyping schemas. The presence of a *k*-mer was used a surrogate for the presence of the corresponding SNP. Each position was assigned one of three states: *ref* (wild type), *alt* (variant), or *no call* (includes missed and ambiguous calls). The reference sequences and Python scripts used to extract the detected variants are provided in the Supplementary Materials.

#### Contamination detection at different coverage cut offs

To assess BioHansel’s ability to identify contamination, one representative dataset was selected for each of BioHansel’s five genotyping schemas from each of the 13 most abundant and fully resolved BioHansel genotypes present in the data deposited in the NCBI SRA as of December 2018. Unlike the concordance study above, the genotypes selected for this study included closely related lineages differentiated by very few target SNPs, representing the most likely mixed pairs that might be encountered when sequencing of these pathogens. Accessions are provided in Supplementary Tables S7–S11.

Genome assemblies were generated using Unicycler version 0.4.7 using default settings. Synthetic reads were generated using ART Illumina v. 3.11.14 (Huang *et al.*, 2012) to generate from each of these sequences a synthetic paired-end Illumina read dataset with 500X coverage. We used the *sample* command from Seqtk v. 1.3 (Li, 2019) with a fixed seed of 42 to randomly subsample the reads from these datasets in various proportions to generate our artificial contamination datasets. For each of the five schemas, 156 mixed datasets were generated (every possible pairing of the thirteen most abundant genotypes). For each mixed pair, a set of reads was generated with simulated contamination levels of 0%, 1.67% (2X), 5% (6X), 10% (12X), 13.33% (16X), 16.67% (20X), 33.33% (40X), and 50% (60X), with a final target genome coverage of 120X for the mixed pair. The scripts used to create the artificial contamination datasets are available in the Supplementary Materials.

We used BioHansel v 2.2.0 to genotype all the artificial contamination datasets with 3X, 6X, and 8X minimum coverage cut offs. The percent *fail* reported by BioHansel was interpreted as equivalent to the percentage of artificial contamination datasets successfully detected by the BioHansel QA/QC module (*e.g.*, 90% *fail* = 90% success at detecting contamination).

### BioHansel compute performance

We benchmarked BioHansel for computing speed and memory usage. We used 1,017 *S*. Enteritidis WGS datasets listed in Supplementary Table S12. Additional benchmarking was conducted using 1,000 publicly available read datasets for each of the *Salmonella* serovars (SE, SH, ST, and Typhi). Reads were selected and assembled as described above (see *Salmonella* Typhi WGS dataset construction). Accessions for all datasets are provided in Supplementary Tables S13–S16. We used the reference sequences used to generate the genotyping schemas (accessions are provided in the Supplementary Materials) as templates to generate synthetic sequencing reads. Synthetic paired-end Illumina read datasets were generated with ART Illumina v. 3.11.14 (Huang *et al.*, 2012) from each of these sequences using a fixed seed of 42. The final coverage level was set at 10X, 50X, 100X, and 1,000X.

BioHansel v 2.2.0 was benchmarked for runtime and memory against Snippy v. 4.3. Both BioHansel and Snippy were run on unassembled WGS reads using a Linux workstation running Ubuntu 16.04.6 LTS with dual-socket 2.00 GHz Intel(R) Xeon(R) E5-2660 v. 4 CPUs (14 cores per socket), 256 GB RAM, and a Samsung SSD 860 EVO 4 TB hard drive. The benchmarking procedure is available as a NextFlow workflow in the Supplementary Materials. BioHansel was designed to run in parallel so it can be deployed in high performance compute clusters, but can also be run on a regular laptop. To demonstrate this, we ran BioHansel on a modern laptop running Ubuntu 18.04.2 LTS x86_64 with a 3.20 GHz Intel i5-6500 (4) CPU and16 GB RAM. Additional benchmarking results are provided in the Supplementary Materials.

## Results and Discussion

Traditional SNP-based phylogenomics workflows are highly discriminatory, accurate, and reproducible, but are computationally intensive, must be recomputed to incorporate newly sequenced isolates, and require expert knowledge to generate and interpret. These limitations have impeded their uptake by health authorities for infectious disease surveillance activities. Gene-by-gene methods, which can also be expensive to initially compute, need typically be computed only once; they also report their results as simple genotypes, which are familiar to epidemiologists, microbiologists, and the broader health profession. For these reasons, gene-by-gene methods serve as a ready drop-in replacement for existing infectious disease diagnostic and surveillance programs. Gene-by-gene methods were among the first WGS methods to be operationalized for infectious disease surveillance, most notably by the PulseNet International network for foodborne disease surveillance, which has largely replaced its traditional pulsed-field gel electrophoresis-based molecular surveillance with core-genome and whole-genome MLST (Nadon *et al.*, 2017). BioHansel is a hybrid of the SNP and gene-by-gene approaches: it employs a fast *k*-mer matching algorithm to identify diagnostically informative SNPs in a hierarchically structured schema to rapidly and efficiency genotype isolates from raw or assembled WGS datasets, allowing it to accurately place an isolate within a pathogen’s population structure. Like gene-by-gene methods, BioHansel relies on a predefined schema in order to assign a genotype. The principal advantages of BioHansel over gene-by-gene methods are its speed, its minimal computing requirement, its simplicity, its ability to work with both raw reads and assembled contigs, the stability of its schema design, and its inherent ability to identify intra-species contamination.

While tools and pipelines exist that can assign lineages from SNP data, such as Genotyphi (Holt, 2019; Wong *et al.*, 2016), TB-Profiler (Coll *et al.*, 2015) and MTBseq (Kohl *et al.*, 2018), they are developed for—and are tied to—the specific pathogen schemas that they use to call SNPs and assign genotypes, whereas BioHansel is designed to genotype any pathogen that has an appropriately constructed SNP schema. Existing tools also lack unambiguous determination of mixed strains. Another major advantage of BioHansel over these programs is the ability to genotype directly from WGS reads, which avoids costly genome assembly and performs better quality assurance and quality control by taking into account the SNP coverage present in the sequence read data.

### BioHansel quickly and accurately genotypes isolates from WGS data

We designed BioHansel to rapidly and accurately genotype bacterial pathogens using a hierarchically structured SNP-based genotyping schema and nomenclature. To demonstrate BioHansel’s genotyping ability, we used 59 sporadic and outbreak-associated *S*. Heidelberg sequences that were previously analysed using SNVPhyl, a traditional SNP-based phylogenomic pipeline that has been validated for the analysis of foodborne disease pathogens (Bekal *et al.*, 2016). In addition to performing the whole-genome SNP comparison, we also compared the isolates by wgMLST using the BioNumerics software platform and the *Salmonella* wgMLST schema used by PulseNet Canada for outbreak detection. The results are presented in Figure 2 and in the Supplementary Materials.

**Figure 2.**
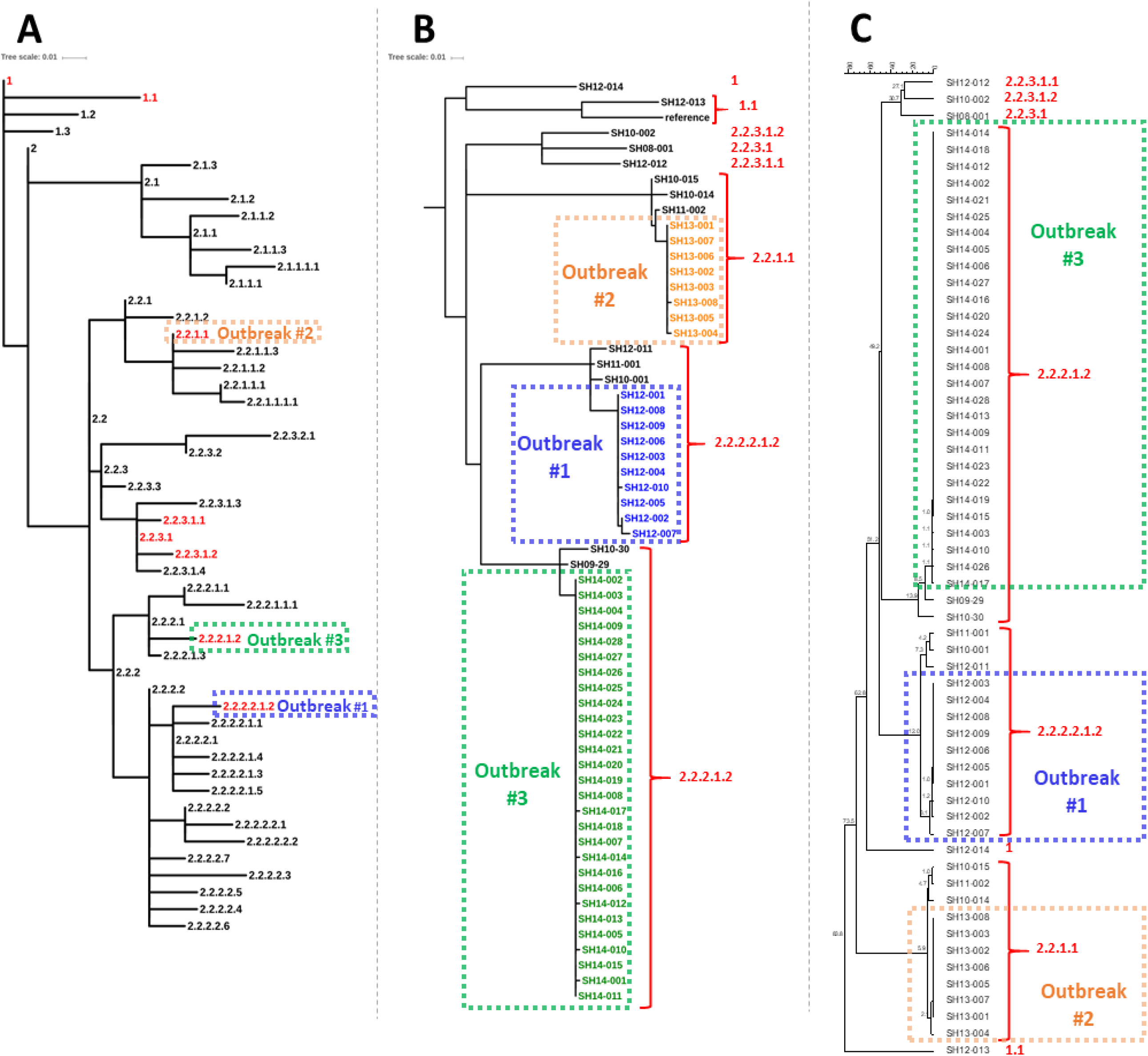
Analysis of retrospective *S*. Heidelberg outbreaks and sporadic and historical cases from 2008 to 2014 in Quebec, Canada. The BioHansel genotypes for the retrospective dataset strains are shown in red font in all panels. **Panel A**: Backbone of the population structure of *S*. Heidelberg showing the genotypes defined in the SH schema v. 0.5.0. The tree was generated the 202 canonical SNPs used in the SH schema (see Supplementary Table S20). The scale bar is equivalent to a distance of ~2 SNPs. **Panel B**: Phylogenetic tree produced by the SNVPhyl pipeline (Petkau *et al.*, 2017) for the 59 *S*. Heidelberg isolates. The scale bar is equivalent to a distance of ~3 SNPs. **Panel C**: UPGMA tree from wgMLST analysis generated using BioNumerics v. 7.6.3 on 4,396 alleles. The scale bar at the top indicates the allelic distance (0–80 alleles).

BioHansel classified the three outbreaks as three distinct genotypes. For all three outbreaks, BioHansel correctly grouped the outbreak strains together, but did include some closely related sporadic strains differing from the outbreak strains by fewer than 15 SNPs (Figure 2B, Supplementary Table S21) or 14 alleles (Figure 2C). BioHansel rapidly classifies the isolates and allows users to place the outbreak isolates in the context of the whole population (Figure 2A). For example, it is difficult to assess how closely related the tree branches are in Figures 2B and 2C without the context provided by the pathogen’s broader population structure. Comparing the BioHansel genotype assignments with the whole *S*. Heidelberg population structure (Figure 2A) shows that the isolates in genotype 1 and those in genotype 2.2.2.2.1.2 in Figures 2B and 2C are separated by a genomic distance that practically spans the whole diversity of *S*. Heidelberg and thus are relatively distantly related.

### BioHansel’s genotyping results have high concordance with traditional SNP calling workflows

To evaluate BioHansel’s SNP-detection accuracy, we compared its concordance with Snippy (Seemann, 2015), a traditional SNP-detection workflow. We chose to compare BioHansel with Snippy as it is popular, built for speed, and has demonstrated accuracy that compares favourably to other traditional SNP-detection workflows (Whaley *et al.*, 2018).

Unlike traditional SNP genotyping workflows that identify SNPs by mapping reads to a reference, BioHansel uses a fast, exact *k*-mer matching algorithm to search reads or assembled contigs for a predefined set canonical SNPs targeted by *k*-mers contained in a hierarchically structured schema. BioHansel computes the number of times an exact match is found for each of the *k*-mers in the schema and reports an average *k*-mer coverage based on the number of times each *k*-mer target was found. As BioHansel only considers the SNPs targeted by the *k*-mers contained within a given schema, we used an existing *S.* Typhi SNP schema developed by Wong *et al.* for our concordance assessments (Wong *et al.*, 2016). The dataset consisted of 1,000 high-quality sequences drawn from the EnteroBase, a publicly available database of enteric pathogen genomes (Alikhan *et al.*, 2018). BioHansel average *k*-mer coverage correlated well with the Snippy mean read mapping coverage (*r*_Pearson_= 0.99), but was approximately 73% of the mean mapping read coverage (Figure 3 and Supplementary Table S22). When comparing the read mapping coverage obtained using Snippy at each target SNP position, the average BioHansel *k*-mer coverage also showed consistent correlation with the Snippy mapping coverage with very few exceptions (*r*_Pearson_= 0.97, Figure 3B). The lower *k*-mer coverage estimate results from BioHansel’s detection algorithm, which identifies only exact matches to the target *k*-mers. BioHansel’s ability to make an exact *k*-mer match depends on the size of the target *k*-mers, the genomic diversity in the supplied sequence data (*i.e.*, the presence of additional mutations within the genome sequences targeted by the *k*-mers), its GC content, and the level of sequencing error.

**Figure 3.**
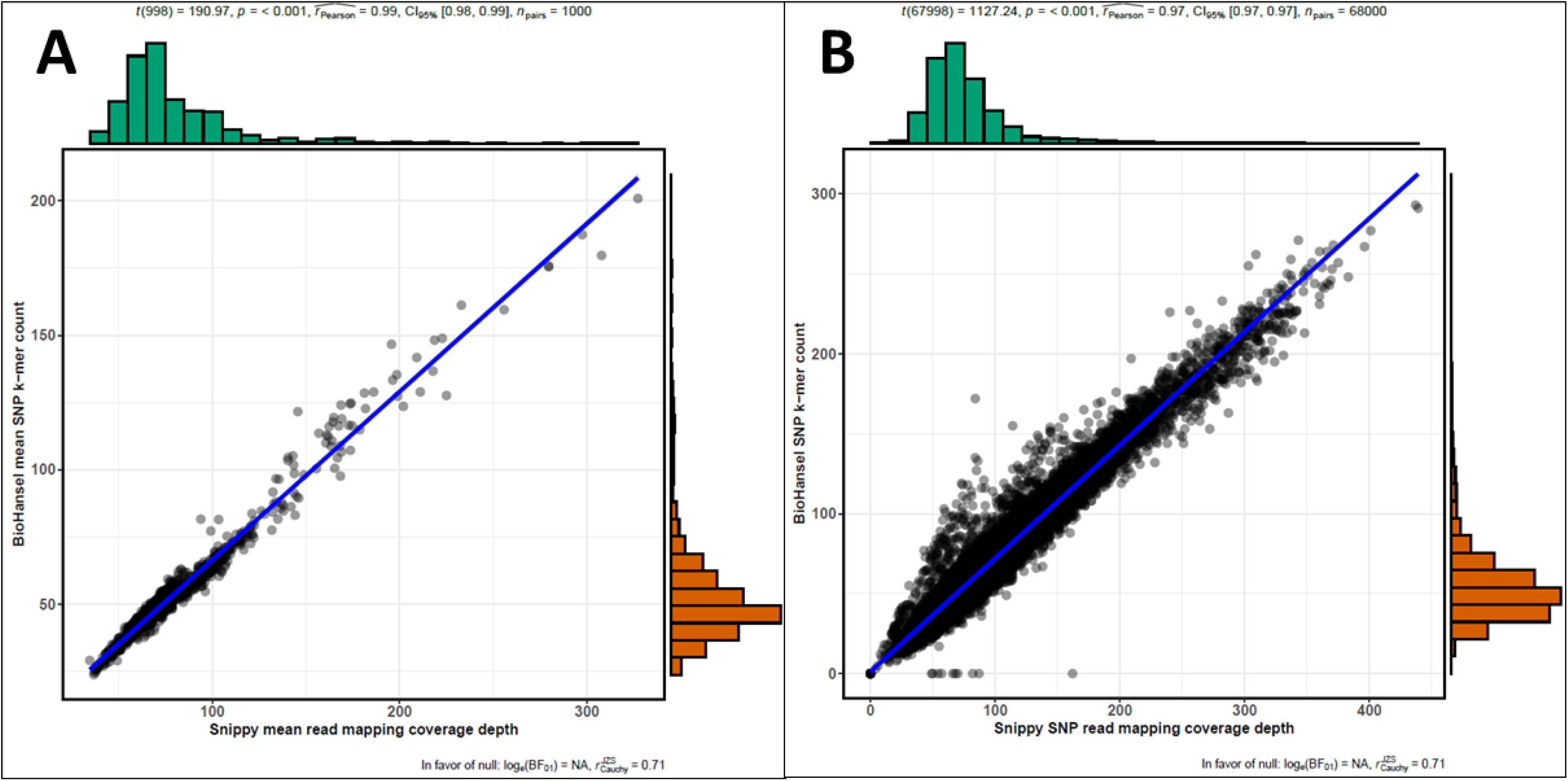
SNP coverage correlations between BioHansel and Snippy. **A.** Scatterplot of the average BioHansel *k*-mer coverage using the Typhi schema with 68 SNP targets, against the mean read mapping coverage of Snippy for 1,000 public *S.* Typhi genomes. **B.** Scatterplot of the BioHansel *k*-mer coverage against the Snippy read mapping coverage at the 68 SNPs targeted by the Typhi genotyping scheme (*n* = 1,000 genomes). The blue lines represents the line of best fit. SNP coverages calculated by BioHansel are consistently around 73% of the Snippy sequence mapping coverage.

The genotyping results obtained by BioHansel were compared with those of Snippy at 3X, 6X, and 8X minimum coverage thresholds (Table 1 and Supplementary Tables S23–24). Both tools detected the same base > 99.9% of the time for the genotyping SNP sites with zero cases of discordant majority bases calls (Table 1), demonstrating that BioHansel produces results that are highly concordant with traditional SNP-calling methods.

For all three of the coverage thresholds evaluated, Snippy was able to detect slightly more bases than BioHansel. The most substantial differences were observed at the 3X coverage cut off (Table 1). (At this coverage level, a target *k*-mer need only be present in three reads in order meet the coverage threshold.) Both tools identified both inclusion and exclusion *k*-mer targets for some of the SNP sites defined in the Typhi schema; these were considered mixed calls. There are cases where BioHansel was able to identify a base unambiguously, whereas Snippy reported a mixed call, and *vice-versa* (Table 1). This discrepancy is likely due to some degree of low-level contamination in the WGS datasets, systematic sequencing errors, or random sequencing errors in areas of high coverage. There are very few differences between the tools at the higher cut off thresholds (Table 1). Notably, the number of mixed calls by both tools drops sharply as the minimum coverage threshold is increased from 3X to 8X, while the number of missing bases stays relatively constant. These results demonstrate that using the BioHansel default minimum coverage of 8X effectively eliminates the noise from the results without unduly compromising SNP detection at the total sequence coverage levels typically generated for these pathogens, and that the BioHansel SNP calls are in excellent agreement with those of Snippy at this threshold.

We also compared our BioHansel genotyping results with the results from the Genotyphi companion tool for *S.* Typhi genotyping that, like BioHansel, uses a hierarchical SNP schema (Holt, 2016). Since our Typhi schema is adapted from this work, we are able to perform a straightforward comparison. We tested the same 1,910 isolates (Supplementary Table S25) used in the original Genotyphi publication (Wong *et al.*, 2016). Of these, 1,894 (99.16%) passed BioHansel’s QC checks while 16 failed QC (Supplementary Table S26). Both BioHansel and Genotyphi generated identical genotypes for all isolates that passed QC. For the samples failing QC, BioHansel’s QC module provides us with some insight into why this may be the case. One of the isolates was not a true Typhi but the outgroup strain Paratyphi A. another isolate was missing a target defining an intermediate hierarchy level in the final subtype (poor quality, low coverage). Three isolates gave mixed results and were missing > 5% of the *k*-mer targets for the contaminant genotype. Considering that these three isolates had > 150X average *k*-mer coverage, it is hard to tell if this is noise or real contamination, as the detected contaminant genotypes are closely related to the dominant genotypes. The other 11 isolates that failed BioHansel QC gave mixed results; of these, 2 appear genuinely mixed as multiple targets were detected that support 2 genotypes. For the other 9, only one “contaminant” *k*-mer target was detected, likely due to excessive noise detected at the 8X QC threshold. (Notably, 4 out of the 9 isolates had an average *k*-mer coverage above 100X.) A detailed analysis of these BioHansel results is provided in the Supplementary Materials. These results showing that even in the samples that failed QC, *k*-mers matching the *S*. Typhi genotype previously identified by Genotyphi were also detected by BioHansel. Based on these results, we conclude that BioHansel successfully identified all genotyping targets using the adapted Typhi schema and, compared to Genotyphi, provides additional QC information about *k*-mer target abundances when analysing read data.

### BioHansel detects intra-species contamination

BioHansel’s hierarchically structured schemas lend themselves naturally to the detection of intra-species contamination, a novel functionality not provided by other SNP-based genotyping tools. BioHansel identifies contamination in WGS datasets in two ways: 1) detection of heterozygous SNPs and 2) the presence of SNPs with conflicting genotype assignments. We assessed the ability of the BioHansel QC module to identify contamination in artificially constructed WGS datasets by mixing two distantly related genotypes of the same pathogen together in different proportions. We also examined how the minimum coverage threshold for SNP detection affected BioHansel’s ability to detect contamination.

In the first experiment, the effect of contamination on the detection of SNPs was examined by running Snippy and BioHansel on artificial contamination datasets generated using the most divergent genotypes for each of the five schemes currently implemented in BioHansel. The comparative analysis between the results from the two tools was restricted to the SNP positions used by BioHansel for genotyping each pathogen. By mixing synthetic WGS data of isolates from different BioHansel genotypes, it is expected that base-calling conflicts will occur at one or more of these target SNP positions for each of the artificial contamination datasets. The selected strains are listed Supplementary Table S5 and schema-specific results are available in the Supplementary Tables S27–S29. A graph showing the results obtained with the artificial contamination datasets at 15 different contamination ratios for all five schemes is presented in Figure S3. Both tools reported unambiguous base calls in 92.85% of the 16,185 target SNP bases considered across all contamination levels (Supplementary Table S30). There were no cases where both tools reported unambiguous, but conflicting variants. As expected, as the coverage of the contaminated genome increased, both tools reported increased ambiguous base calls (Figure S3). Both tools show a similar distribution in calling bases that the other tool did not call (Figure S3). In total, BioHansel unambiguously detected 97 variants where Snippy was ambiguous (0.6% of all base calls), whereas Snippy unambiguously detected 66 where BioHansel was ambiguous (0.4% of all base calls; see supplementary Table S30). Taken together, these results indicate that BioHansel readily detects contamination *k*-mers above the default threshold of 8X depth of coverage in isolates and confirms that differences observed between the tools are the result of ambiguity in identifying variants and not from the detection of conflicting variants.

In a second experiment, we mixed publicly available sequence data for the 13 top genotypes identified for each of the 5 pathogens, regardless of the evolutionary distance between them, in order to replicate the types of contamination that a sequencing laboratory might encounter. The results presented in Figure 4 show that BioHansel successfully detected all contaminated datasets at all coverage thresholds tested when the contamination level is above 16% (Figure 4 and Supplementary Tables S31–S35). The results also show that the tool’s ability to detect low levels of contamination (< 10%) increases with the use of lower coverage thresholds, however, this comes at a cost. When using the lowest *k*-mer coverage threshold of 3X (*i.e.*, the *k*-mer was detected in a minimum of 3 reads), the SH, SE, & Typhi schemas (which use multiple targets to assign a genotype) reported contamination when no contamination was present (Figure 4 and Supplementary Tables S31–S32; S34). Under these conditions (i.e. using 3X coverage threshold), between ~8%–23% of the uncontaminated datasets falsely identified sequence error as contamination, in contrast to the results obtained using 6X or 8X thresholds where the sequence error was below these coverage thresholds and did not trigger a false contamination QC error in the uncontaminated datasets. We conclude that the higher coverage cut offs at or above 6X eliminate the “background noise” caused by the presence of sequencing errors in uncontaminated datasets, but will also mask low levels of contamination (< 10%). We used these findings to set BioHansel’s default values, which are intended for high quality Illumina WGS data and for assembled contigs, but may need to adjusted for users with custom schemas or when analyzing sequence data with high error rates, such as long-read sequence data.

**Figure 4.**
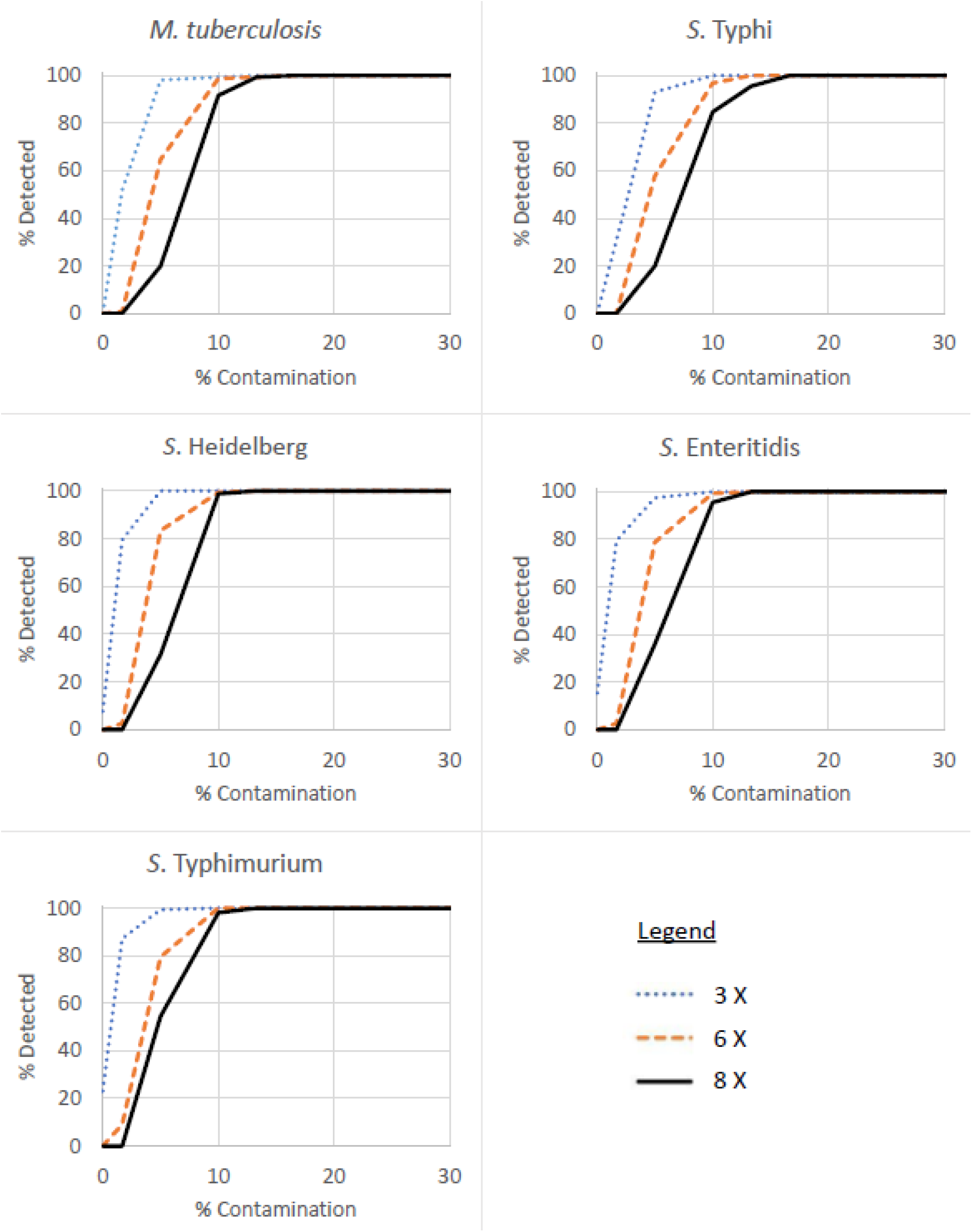
Contamination Detection Performance. 156 contamination datasets were constructed for each pathogen. Each graph represents the percentage of contamination datasets detected by BioHansel (Y axis) among all the artificially generated datasets tested for each contamination levels (X axis). The minimum SNP detection thresholds tested are shown in the legend.

BioHansel’s ability to detect contamination in WGS datasets also improves with increasing numbers of *k*-mer targets used to assign a genotype (Figure 4). The MTB and Typhi schemas, which use only a single canonical SNP to assign a genotype, require slightly higher levels of contamination to be detected by BioHansel (Figure 4). This is particularly evident from our analysis of data sets with low levels of contamination, since the schemas that performed better at these levels had a larger number of targets and more redundancy for those genotypes. This makes intuitive sense, since at low levels of contamination fewer target *k*-mers will be detected above the minimum read coverage threshold.

#### Interpreting BioHansel’s QA/QC reports

BioHansel generates a QA/QC report that can be straightforwardly interpreted by non-experts for both the genotyping results and the detection of intra-species contamination. When performing whole genome assembly or genome mapping, intra-species contamination manifests as an increase in ambiguity of some SNP positions. Interpreting these results as contamination from a VCF output file requires time and expertise, which is not readily or widely available. As demonstrated, the ability of BioHansel to detect low levels of contamination in unassembled WGS datasets is related to the number of *k*-mers used to define each genotype: the higher the number of *k*-mer targets included in the schema, the more likely they are to generate informative contamination signal from conflicting genotyping results. The interpretation of these results is performed automatically by BioHansel’s QA/QC module and reported for each dataset.

### BioHansel’s speed and memory performance compare favourably to traditional SNP-calling pipelines

#### Runtime and memory usage for genotyping from assembled contigs

The Aho–Corasick algorithm maps *k*-mers to genomic data in linear time. We confirmed this relationship using four of our five schemas (two schemas had approximately the same number of *k*-mer targets). Assembled genomes were genotyped by BioHansel in batches of 1,000 using a single CPU core for each genotyping schema. We observed an average runtime of between 6.8–8.5 minutes, corresponding to between 0.41–0.51 seconds per assembly (Supplementary Figure S4). The runtime increased linearly with the number of *k*-mer targets in the schema (Supplementary Figure S4). Similarly, the memory usage also increased linearly with the number of *k*-mer targets in the genotyping schema (Supplementary Figure S5). When run on a personal computer using only a single CPU core (*i.e.*, one thread) BioHansel runtime averaged 0.23–0.31 seconds per assembly (Supplementary Figure S6). These results highlight the extreme rapidity in which BioHansel can genotype assembled genomes, using minimal computing resources—schemas could easily extend to thousands of targets without appreciably affecting genotyping performance.

#### Runtime and memory usage for genotyping from read data

One of the major strengths of BioHansel is that it can genotype pathogens from both assembled contigs and raw sequence reads. We evaluated the performance of BioHansel using Illumina sequencing reads as input and found that the runtime increased linearly according to the number of *k*-mer targets included in the schema (Figure 5, Supplementary Table S36). The runtime for BioHansel scaled well with increasing sequence coverage of the isolates (Figure 6). These results highlight that within the normal sequence coverage levels for bacterial WGS datasets, BioHansel provides extremely rapid genotyping information.

**Figure 5.**
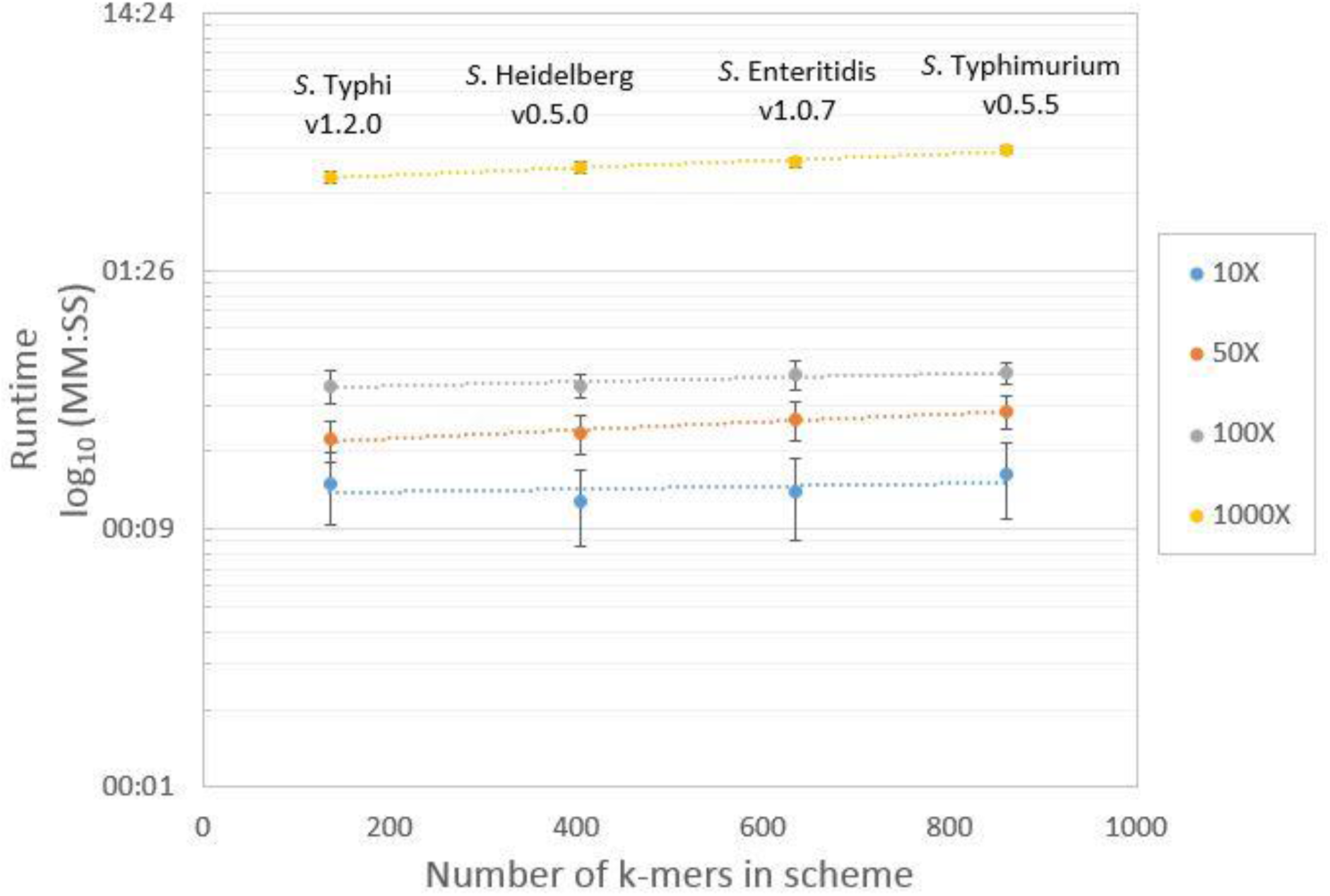
BioHansel runtime for one synthetic Illumina Miseq dataset with increasing genome coverage and numbers of *k*-mer targets. Each data point represents the average BioHansel runtime on 10 different datasets of SH, Typhi, SE or ST, at the genome coverages depicted.

**Figure 6.**
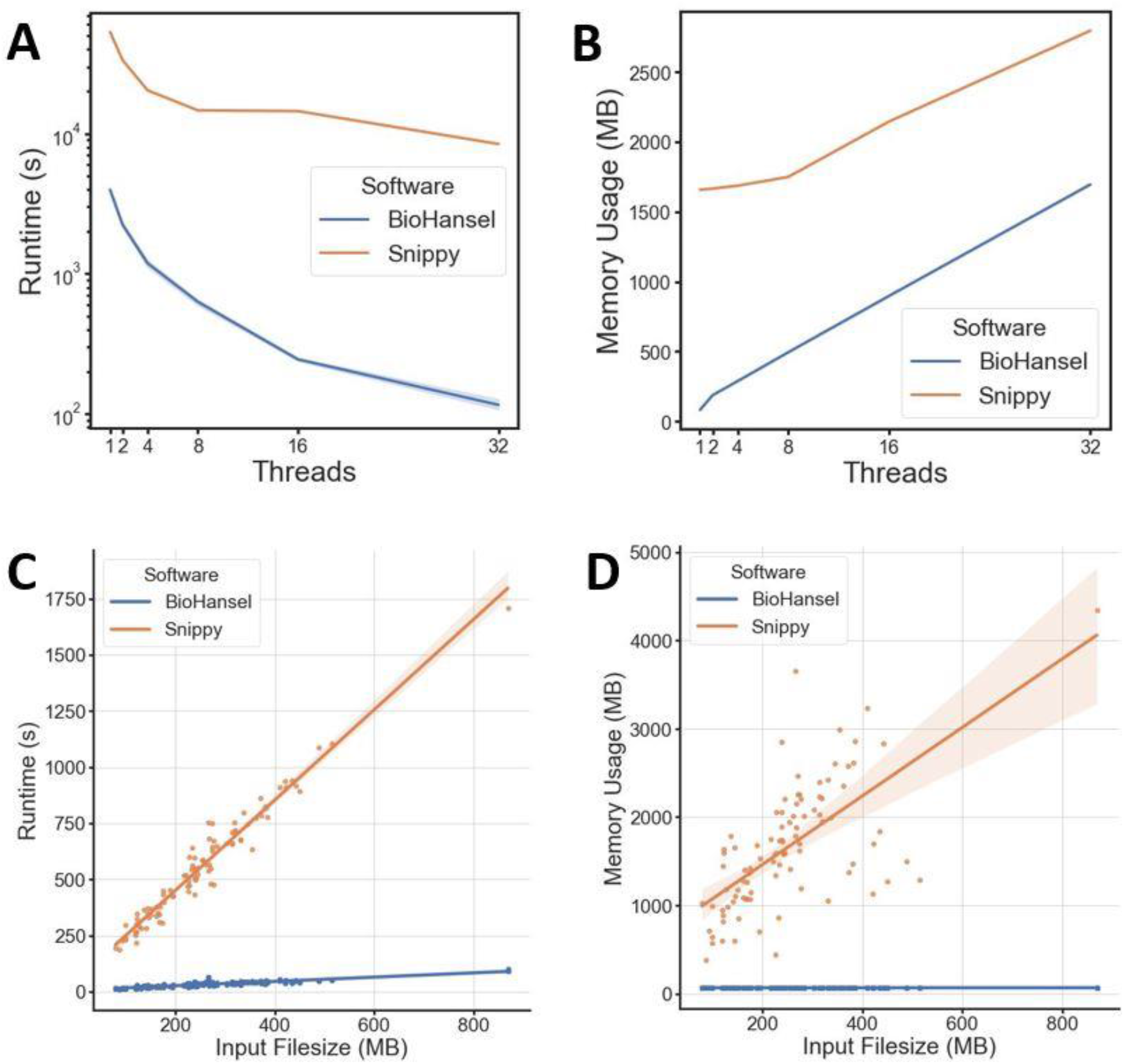
Runtime and memory usage comparison between BioHansel and Snippy for unassembled Illumina WGS datasets from *S*. Enteritidis. **Panels A and B:** Runtime and memory usage for 96 WGS datasets with increasing numbers of parallelized compute threads. **Panels C and D:** Runtime and memory usage for single unassembled WGS datasets of increasing size (*n* = 96) using a single CPU core. The shading in the graphs represents the 95% confidence interval with 1,000 bootstraps using a mean aggregation estimation.

To provide context on BioHansel’s speed, its runtime was compared to Snippy, a traditional SNP-calling workflow. Since Snippy is designed for speed (Seemann 2015) it should provide a lower bound estimate of runtimes for traditional SNP-calling workflows. As with most SNP calling workflows, Snippy reports all the SNPs detected in the genome relative to a given reference strain, whereas BioHansel considers only the SNPs required for genotyping. BioHansel and Snippy were used to analyze artificially constructed datasets at different levels of sequence coverage. BioHansel’s runtime at 10X coverage ranged from 11–14 seconds compared to 65–72 seconds for Snippy, which represents a nearly six-fold reduction in runtime (Supplementary Table S35). Further gains were observed increasing sequence coverage levels with approximately a fourteen-fold reduction in compute time at 50X coverage. With a recommended minimum genome coverage of 60X to detect all the SNPs in a bacterial genome (Kozyreva *et al.*, 2017), this represents a substantial time savings.

To exemplify the difference in runtime and memory usage between the tools in a typical public health laboratory setting, we have compared the tools using 96 public datasets of *S*. Enteritidis on a personal laptop computer (Figure 6). The results show that the BioHansel runtime is over an order of magnitude lower than that of Snippy. Increasing the number of threads (cores) significantly reduced runtime for both tools (Figure 6A), while the memory usage increased linearly for both tools (Figure 6B). When comparing the analysis of individual datasets, we observed that the file size, a proxy for the number of reads, does not have as much of an effect on BioHansel’s runtime as it does on Snippy, which increased sharply with file size (Figure 6C). Snippy’s memory usage also increased with file size while BioHansel’s memory usage remained constant (Figure 6D). We note the runtimes recorded for Snippy do not take into account the time required to build the subsequent phylogenetic tree; given the number of isolates typically required to contextualize an outbreak, and the tree building algorithm used, this process can take hours to days.

## Conclusions

BioHansel was designed to address the need for genotyping pathogens in a fast, flexible, and readily interpretable manner without the need for high performance computing or specialized scientific expertise. We demonstrated that BioHansel rapidly and accurately genotypes *Mycobacterium tuberculosis* and four prevalent subtypes of *Salmonella*. BioHansel’s SNP detection is highly concordant with Snippy, a traditional SNP-calling pipeline, and Genotyphi, a SNP-based genotyping pipeline for *Salmonella* Typhi. We conclude that BioHansel readily identifies SNP targets with high concordance to traditional SNP-calling workflows, while additionally providing phylogenetically informative genotyping codes to typed isolates.

We also demonstrated BioHansel’s ability to leverage read data to identify intra-species contamination. BioHansel’s quality assurance routines can also identify poor quality sequence data as well as sequence data with insufficient read coverage. BioHansel’s use of SNP-targeting *k*-mers increases the specificity of the SNPs detected and vastly reduces the effects of sequence contamination from other organisms (Goig *et al.*, 2018).

We provide schemas for *Mycobacterium tuberculosis* and four prevalent subtypes of *Salmonella*, since these are highly clonal organisms and are amenable to genotyping with hierarchically structured canonical SNP schemas. BioHansel’s ability to genotype less clonal organisms is an open question. BioHansel can genotype with SNPs contained in a pathogen’s accessory genome, although it is not currently configured to accommodate SNP profiles acquired through recombination—this remains a topic of active study in our lab as we work to extend BioHansel’s genotyping approach to other priority pathogens.

BioHansel’s genotyping accuracy is a function of the clonality of the target pathogen and the discriminatory power of its canonical SNPs. Traditional gene-by-gene and SNP-based methods that exploit the full diversity of sequenced pathogen isolates will naturally provide the highest resolution genotypes and phylogenies. BioHansel trades resolution for speed, but as we demonstrate with as little as 68 canonical SNP targets, still retains sufficient resolution to rapidly identify and discriminate outbreak-related strains from all but the most highly related sporadic circulating strains. Thus, BioHansel may be applied to rapidly generate an initial genotype for rapid response while slower, more compute intensive methods work to capture the full genomic diversity and assign a more specific genotype. BioHansel’s demonstrated ability to type WGS data quickly on computing resources as modest as a modern laptop makes it especially well-suited for use in smaller labs and labs in developing nations that might not have ready access to the high performance computing resources required for traditional WGS-based workflows for surveillance and outbreak investigation.

BioHansel is designed to accept custom schemas allowing users to extend existing schemas or develop novel schemas, although manual schema creation process requires considerable expertise and time investment. Schema development requires bioinformatics and organism expertise to ensure the selection of high quality *k*-mer targets that can reliably genotype target pathogens. In order to overcome this limitation, future work will focus on the automation of schema development, updating existing schemas with new genotypes, and the generation of new schemas for other priority pathogens. The ability to accommodate recombination will be explored, and the addition of an alternate workflow geared towards the analysis of metagenomics datasets will also be explored, as the *k*-mer-based strategy used by BioHansel makes it well suited for fast characterization of the pathogen signature sequences in these datasets.

## Author statements

### Authors and contributors

Geneviève Labbé: Conceptualization, Funding acquisition, Methodology, Data curation, Formal analysis, Investigation, Supervision, Project administration, Writing – original draft, Visualization

Peter Kruczkiewicz: Conceptualization, Software, Methodology, Supervision, Formal analysis

Philip Mabon: Supervision, Software, Methodology

James Robertson: Conceptualization, Software, Methodology, Data curation, Formal analysis, Investigation, Funding acquisition, Writing – original draft, Visualization

Justin Schonfeld: Software, Methodology, Writing – original draft

Daniel Kein: Data curation, Formal analysis, Investigation

Marisa A. Rankin: Data curation, Formal analysis, Investigation

Matthew Gopez: Software, Methodology, Formal analysis

Darian Hole: Software, Investigation

David Son: Software, Methodology, Data curation, Investigation

Natalie Knox: Writing – review & editing

Chad R. Laing: Software, Methodology, Resources

Kyrylo Bessonov: Software, Methodology

Eduardo Taboada: Funding acquisition, Supervision, Writing – review & editing

Catherine Yoshida: Supervision, Project administration, Writing – review & editing

Kim Ziebell: Funding acquisition, Formal analysis, Investigation

Anil Nichani: Funding acquisition, Supervision, Writing – review & editing

Roger P. Johnson: Funding acquisition, Project administration, Supervision, Writing – review & editing

Gary Van Domselaar: Funding acquisition, Project administration, Resources, Supervision, Writing – review & editing

John H.E. Nash: Funding acquisition, Project administration, Supervision, Writing – review & editing

### Conflicts of interest

The author(s) declare that there are no conflicts of interest.

### Funding information

This work was supported by grants from the Government of Canada Genomics Research and Development Initiative and the Public Health Agency of Canada.

## Acknowledgements

We thank Jonathan Looi for help with the development of the *S*. Typhimurium subtyping schema and BioHansel Read-the-Docs, Gary Tong for help with the development of the BioHansel Read-the-Docs, Adrian Zetner for help with cluster commands, and Aaron Petkau for help with Galaxy and IRIDA workflows.

## Notes

https://github.com/phac-nml/biohansel-manuscript-supplementary-data

